# TraceSpecks: a software for automated idealization of noisy patch-clamp and imaging data

**DOI:** 10.1101/299875

**Authors:** Syed Islamuddin Shah, Angelo Demuro, Don-On Daniel Mak, Ian Parker, John E. Pearson, Ghanim Ullah

## Abstract

Experimental records of single molecules or ion channels from fluorescence microscopy and patch-clamp electrophysiology often include high-frequency noise and baseline fluctuations that are not generated by the system under investigation and have to be removed. Moreover, multiple channels or conductance levels can be present at a time in the data that need to be quantified to accurately understand the behavior of the system. Manual procedures for removing these fluctuations and extracting conducting states or multiple channels are laborious, prone to subjective bias, and hinder the processing of often very large data-sets. We introduce a maximum likelihood formalism for separating signal from a noisy and drifting background such as fluorescence traces from imaging of elementary Ca^2+^ release events called puffs arising from clusters of channels and patch-clamp recordings of ion channels. Parameters such as the number of open channels or conducting states, noise level, and background signal can all be optimized using the expectation-maximization (EM) algorithm. We implement our algorithm following the Baum-Welch approach to EM in the portable java language with a user-friendly graphical interface and test the algorithm on both synthetic and experimental data from patch-clamp electrophysiology of Ca^2+^ channels and fluorescence microscopy of a cluster of Ca^2+^ channels and Ca^2+^ channels with multiple conductance levels. The resulting software is accurate, fast, and provides detailed information usually not available through manual analysis. Options for visual inspection of the raw and processed data with key parameters, and exporting a range of statistics such as the mean open probabilities, mean open times, mean close times, and dwell time distributions for different number of channels open or conductance levels, amplitude distribution of all opening events, and number of transitions between different number of open channels or conducting levels in asci format with a single click are provided.

## Introduction

Recent advances in total internal reflection fluorescence microscopy (TIRFM) provide a powerful tool for the functional study of thousands of Ca^2+^ release channels simultaneously at a single channel resolution in minimally invasive manner, maintaining the physiological environment of the channels (1–3). This technique, called “optical patch-clamp”, has been used to study the Ca^2+^ flux through several individual channels including N-type voltage-gated Ca^2+^ channels (1, 4, 5), nicotinic acetylcholine receptors (2, 5), and L-type Ca^2+^ channels in cardiac muscle (5). Optical patch-clamp was also used to resolve the quantal substructure of elementary Ca^2+^ puffs and sparklets arising from concerted opening of multiple channels in a cluster of inositol 1,4,5-trisphosphate (IP3) receptors (IP_3_Rs) and dihydropyridine-sensitive voltage-gated Ca_V_1.2 channels respectively, by directly resolving the opening and closing of individual channels in the cluster (6, 7). Recently, the same technique was employed to resolve the conductance levels in a single ion channel while simultaneously imaging thousands of Ca^2+^-permeable plasma membrane (PM) pores formed by beta amyloid (A*β*_1-42_) in the cells with Alzheimer’s disease pathology (8, 9).

Optical patch-clamp generates thousands of time-traces representing the gating of individual channels in a single experiment. However, manual analysis of the data generated is extremely tedious and challenging, which hinders the full utility of this powerful technique. The analysis of fluorescence traces from single channels with a single or multiple conductance levels or concerted opening of multiple channels in a cluster of channels (such as IP_3_Rs, ryanodine receptors, and dihydropyridine-sensitive voltage-gated Ca_V_1.2 channels) obtained from TIRFM requires the removal of background noise and baseline fluctuations in the data records that are not generated by the channels under study. These fluctuations could arise either from the saturation of dye molecules or drift in measuring equipment itself. Since the data is expected to exhibit quantized steps, the signal and the data (and possibly the noise) have significant autocorrelation times, and simple filtering may not yield a clear separation of signal from background and fails to make use of the known quantal character of the data.

A similar situation arises while using other techniques such as single channel patch-clamp and single molecule fluorescence and photobleaching experiments, where analysis of the experimental data first requires the removal of noise and varying background that is not generated by the phenomenon under study (10–15). In the patch-clamp experiments (16), instabilities in the gigaohm seal formed between biological membrane patch and patch-clamp microelectrode often result in substantial fluctuations in the magnitude of the observed current and a drift in the recorded signal. Such baseline fluctuations are usually significantly slower than the abrupt changes in current magnitude caused by opening and closing of the channel, and have to be removed before the current traces can be analyzed further by standardized software such as QUB (17, 18) and HJCFit (19) that model the channel gating characteristics based on idealized data.

Existing baseline subtraction algorithms can require substantial time and/or user interaction on ion channel data as the program forces user to check each individual fit by eye. Without a robust baseline subtraction algorithm, some experimentalists perform this laborious task by hand where the experimenter follows the quantal jumps by eye and subtract the drifting background. Such manual background-subtraction procedure requires eye-balling and mouse clicking to indicate the background trajectory, which is time consuming and susceptible to subjective bias. To overcome these obstacles, we previously developed a minimally-parameterized likelihood (20) approach for separating the signal from noisy and drifting background, which gives results that agree with the mouse-based method and is much faster (21). However, the requirement to compile and execute the algorithm for processing the data proved a major hurdle for the experimentalists not familiar with compiled languages. While the main goal of this study was to extend the algorithm so that it can be used for both patch-clamp electrophysiology and fluorescence data, we develop a user-friendly graphical user interface (GUI) which is easy to use, flexible, and portable. Furthermore, we incorporate several new features in the software that were not included in the previous version. In addition to extracting idealized trace and drifting background, the software extracts many features from the data, including the mean open probabilities (P_*O*_), open times (*τ*_*O*_), close times (*τ*_*C*_), dwell-time distributions for different conductance levels or number of open channels, overall P_*O*_, *τ*_*O*_, and *τ*_*C*_ in the trace, amplitude distribution of all events, and the transition frequencies between different conductance levels or channels open with a few simple steps. We demonstrate the utility of our software by processing patch-clamp data containing single and multiple channels per trace, fluorescence traces from TIRFM representing Ca^2+^ signals generated by concerted opening and closing of several channels in a cluster of IP_3_Rs, and fluorescence traces representing the Ca^2+^ flux through PM pores with multiple conductance levels, formed by A*β*_1-42_ oligomers associated with Alzheimer’s disease pathology.

## Methods

In this section we first outline the main points of the algorithm (detailed derivation can be found in Ref. (21)) followed by details of how we generate synthetic records representing patch-clamp electrophysiology of a single ion channel with varying signal-to-noise ratio (SNR) as well as synthetic fluorescence traces representing the concerted opening and closing of multiple channels in a cluster of IP_3_Rs. A brief description of sample experimental methods and the data used to demonstrate the utility of the software is also given towards the end of this section.

### Theory

We use the Expectation Maximization (EM) algorithm to estimate the number of open channels or conductance levels, which is treated as missing data. The EM algorithm was originally developed by Baum, Welch, *et al* in the 1960’s and formalized by Dempster *et al* (22–25).

Assuming that *n*_*t*_ is the number of channels open or the conductance level in which the channel is gating and *I* is the current passing through the channel at time *t*, the observed signal at time t is given by *d*_*t*_ = *b*_*t*_ + *In*_*t*_ + *σ*_*ξ*_*ξ*_*t*_. Where *ξ*_*t*_ is discrete-time Gaussian white noise with moments < *ξ*_*t*_ >= 0, 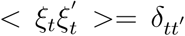. Here *δ*_*tt′*_ is the Kronecker delta and *σ*_*ξ*_ is the noise strength. We further assume that the baseline is undergoing a discrete time random walk, 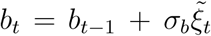 where 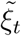 is discrete-time white noise like *ξ*_*t*_. Under these assumptions, it follows that *d*_*t*_ − *b*_*t*_ + *In*_*t*_ and *b*_*t*_ − *b*_*t*−1_ are zero mean Gaussian distributed random variables with variances 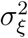 and 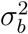 respectively.

The joint distribution for *d, b*, and *n* is given as

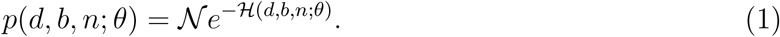

𝓗 is an “energy” function given by:

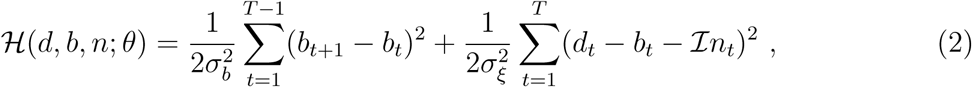

where T is the number of data points, *d* = (*d*_1_,*d*_2_,..…, *d*_*T*_), *b* = (*b*_1_, *b*_2_,..…*b*_*T*_, *n* = (*n*_1_,*n*_2_,..…, *n*_*T*_), and *θ* represents the parameters 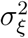, 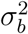, and *I. N* is the normalization factor:

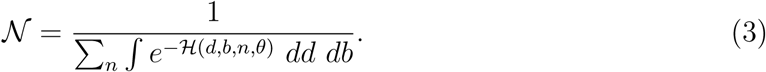

Where 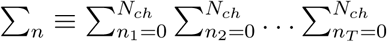 and *N*_*ch*_ is an upper bound on the number of channels contained in the trace (number of channels patched or number of channels in a cluster of channels). For a single ion channel (or molecule) with multiple equally spaced conductance levels, *N*_*ch*_ represents the total number of conductance levels that the system can have, where the closed state of the channel is represented by level 0. *dd* and *db* represent the differentials of all the variables being integrated over: *e.g.* 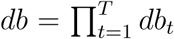. In principal we would like to maximize the likelihood of the data given the model, *L*(*d*; *θ*)

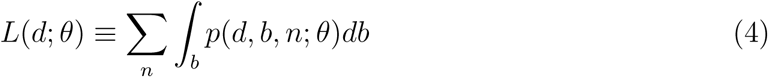

but this is unwieldy. Although the integral over *T* dimensions can be dealt with, the sum is over at least 2^*T*^ values and cannot be maximized in one step. Thus we employ the iterative EM procedure that attempts to find the maximum likelihood estimate of *L* by first making an initial guess for 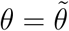 and then iterating the following steps:

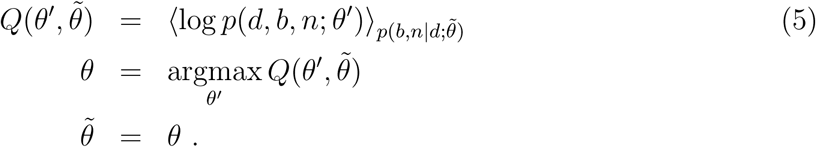

In the above 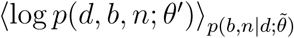 denotes the expectation of log *p*(*d*, *b*, *n*; *θ′*) with respect to the distribution 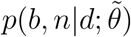 The EM iteration is known to converge to at worst a local maximum likelihood estimate for *L*(*d|θ*).

Because of the simple dependence that the likelihood function has on *b* we can integrate *b* out and then employ a slightly modified version of the EM algorithm to solve the remaining maximum likelihood problem (see Ref. (21) for the derivation).

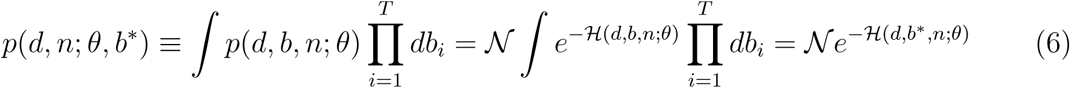

where *b*\* is the maximum likelihood estimate of *b* found by minimizing *H*:

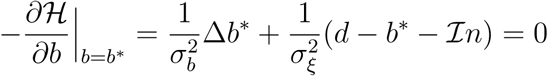

and Δ is the finite difference Laplacian: Δ*b*_*i*_ ≡ *b*_*i*+1_ + *b*_*i*−1_ − 2*b*_*i*_. Thus

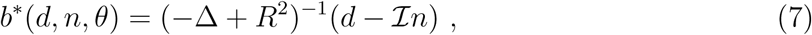

where 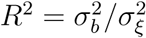 As written, the *b*\* in the argument of *p*(*d, n; θ, b\**) in Eq (6) is redundant since *b*\* is a known function of *d, n*, and *θ*. The reason we have employed this seemingly redundant notation is just to make clear in the algorithm which value of *b*\* (old or updated) we used to compute the current expected values of *n*.

The normalization constant 𝓝 is given by (see Ref. (21) for the derivation):

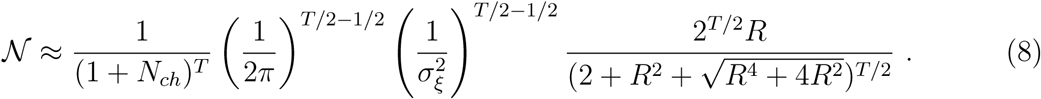

The distribution 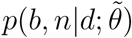 needed for *Q* in Eq (5), becomes 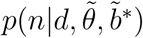 after integrating out *b*. Here 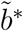 is the initial guess for the background signal. To find 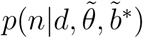 we note that in general *p*(*x*|*y*) = *p*(*x,y*)/*p*(*y*). Thus:

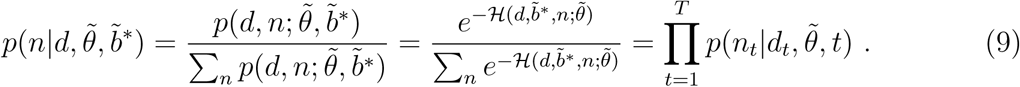

The quantity 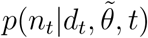 is the conditional probability that there are *n*_*t*_ channels open at the single time point *t*. To simplify the notation, we write 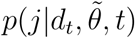 to indicate the conditional probability that there are *j* channels open at time *t*.

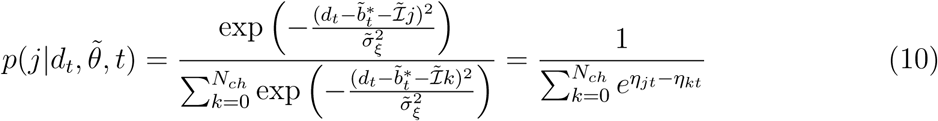

where 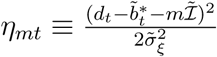, *m* = 0,1,2,3,…*N*_*ch*_. We will use angle brackets without subscripts to denote expectations with respect to 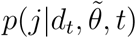 so that

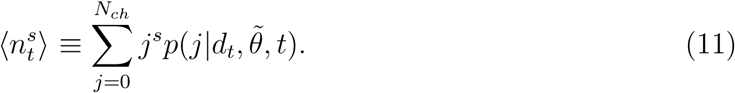

For a single channel or single molecule recording, *N*_ch_ = 1 and Eq (11) results in 〈*n*_*t*_〉 = 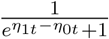

At this point we can write Q explicitly:

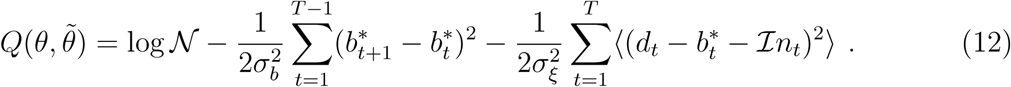

The dependence on 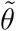 is only through *b*\* and the angle brackets. The end result of this calculation is that we replace the dependence in 𝓗 on powers of *n*_*t*_ with the expected value of those powers: *n*_*t*_ → 〈*n*_*t*_〉 and 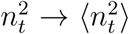 We maximize 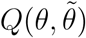 by updating *θ* iteratively in the following steps.

**Step 0:** (Initialization) Make initial guess for 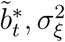, *I*, and 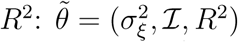

**Step I:** (Estimate or Expectation) Update the expected values of *n*_*t*_ and 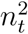 with respect to *p*(*j*|*d*_*t*_, *θ*, *t*) which we will denote: 〈*n*_*t*_〉 and 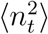.

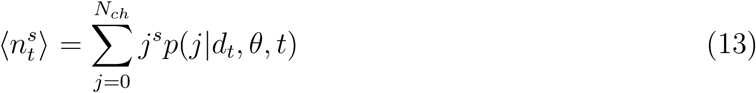

At the end of this step *n*_*t*_ and 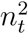 in 𝓗 will be replaced by 〈*n*_*t*_〉 and 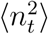 respectively.

**Step II:** (Maximize)

**A. Update baseline signal**

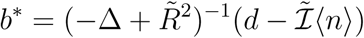

We obtain *b*\* with a standard tridiagonal matrix solver which is very fast* Note the dependence in *b*\* is on 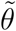 rather than on *θ*. In steps B, C, and D we update the parameters *I*, 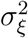 and *R*^2^ while keeping *b*\* fixed.

**B. Update 𝓘.**

The derivative of *Q* with respect to 𝓘 is:

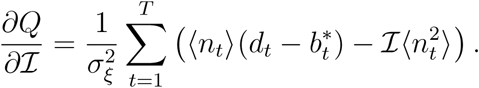

To maximize *Q* with respect to 𝓘 we set the above equation to zero which yields:

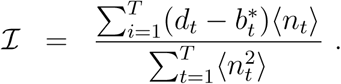

**C. Update** 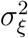

The derivative of *Q* with respect to 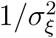 is:

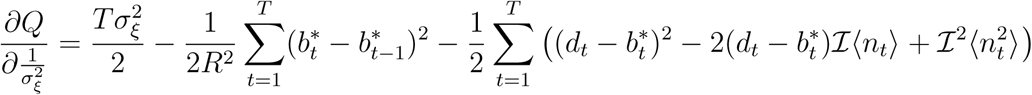

To maximize *Q* with respect to 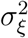 we set the above equation to zero which yields:

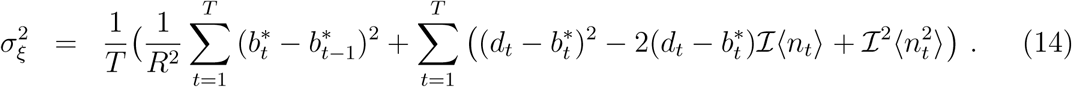

**D. Update *R*^2^**

The derivative of *Q* with respect to *R*^2^ is: 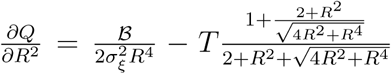 where 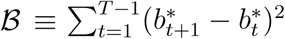 The updated *R*^2^ is the single real positive root of 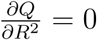. At this point 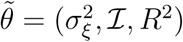.

**Return to Step I.**

To summarize, we iterate the steps described above in the order **Step 0 → Step I → Step II → Step I**. So, in the E step, we compute the *N*_*ch*_ expectation values of *n*_*t*_ for each *t* and in the M step, we update the parameters based on maximizing *Q* in Eq. 6.

The algorithm is implemented in widely portable java language in a user-friendly GUI with options to setup initial parameters and graphical display of raw and processed traces. Options to store statistics on the processed data including mean P_*O*_, *τ*_*O*_, *τ*_*C*_, dwell-time distributions for different conductance levels or number of open channels for the processed trace are provided. Information regarding the amplitude distribution of all opening events as well as the frequency of transitions between different number of open channels or conductance levels are also provided. An example of algorithms GUI is shown in Fig 1. A compiled and portable java program with the GUI that performs these calculations and user manual will be provided in the online supplement after the paper is accepted for publication.

**Fig. 1:**
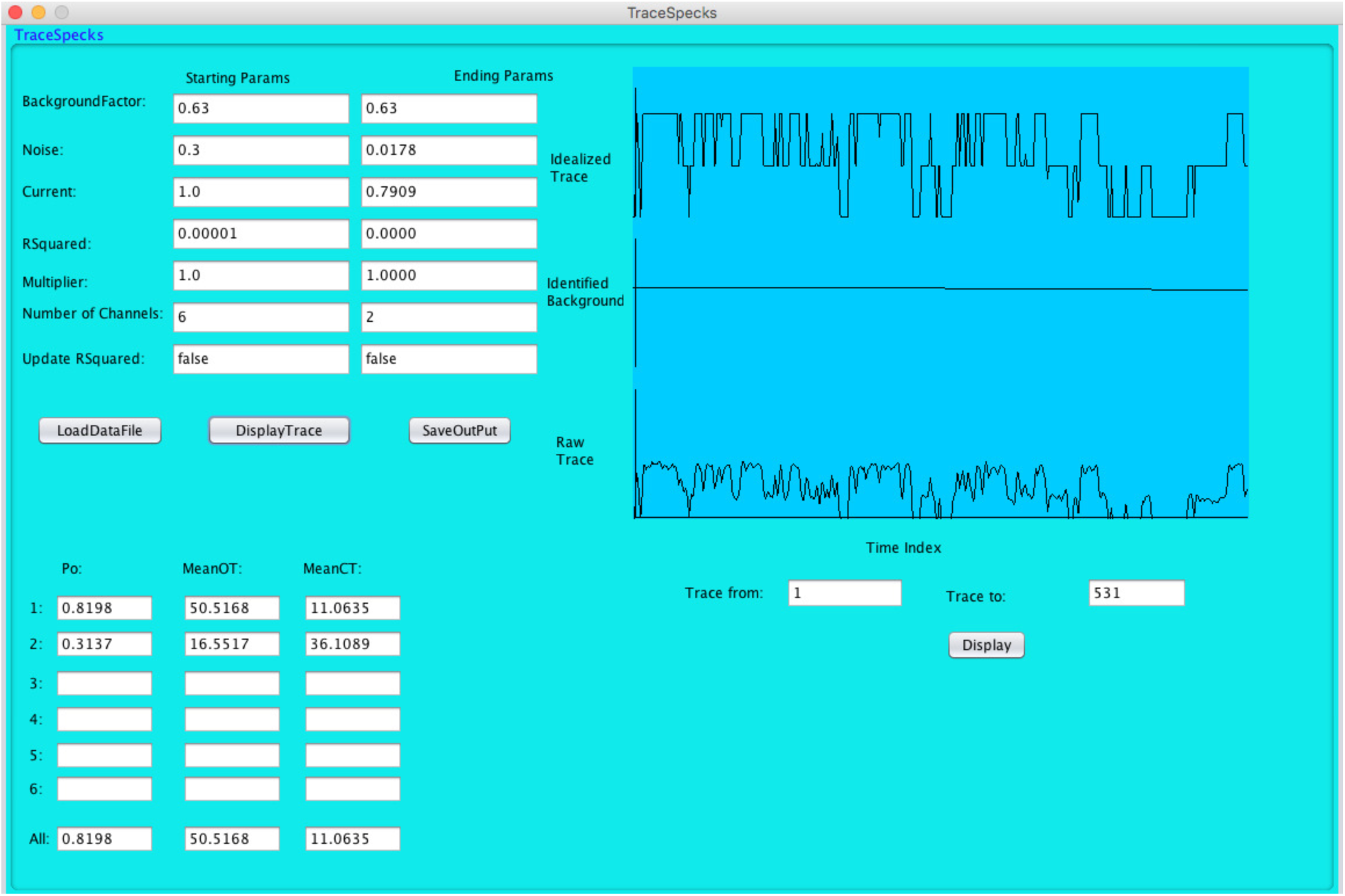
Graphical user interface of TraceSpecks. Various controls for reading, displaying, and saving data sets are provided. Text controls are used to provide input parameters for the algorithm as well as displaying the optimum parameters and results determined by the algorithm. Raw trace, background noise signal, and idealized signals can be viewed in the graphical panel of the GUI for visual inspection and can be zoomed in and out as needed.

### Data Records

We apply our method to synthetic and experimental data sets from both TIRFM experiments and patch-clamp recordings. Our synthetic data sets comprise of two examples, one representing patch-clamp experiments with variable SNR and another one mimicking florescence traces from a cluster of IP_3_Rs (26). Experimental data set consists of nuclear patch-clamp recordings from IP_3_ in Sf9 cells (27), florescence recordings from IP_3_-induced Ca^2+^ puffs (28), and Α*β*-induced Ca^2+^-permeable PM pores (8) in Xenopus oocytes.

#### Synthetic data sets

Details of patch-clamp synthetic data generation can be found in our previous article (21). Briefly, we use 10,000 open and closed events with duration of these events drawn randomly from a uniform distribution. The range of distribution for close events is 5 times that of open events. Channel’s P_*O*_ is controlled by varying the range of the distribution from which the close time is drawn. We construct background signal 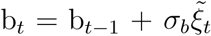 to get final simulated channel trace d_*t*_ = b_*t*_ + 𝓘 n_*t*_ + *σ*_ξ_*ξ*_*t*_, where 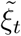 and *ξ*_*t*_ are zero mean Gaussian noise with deviation *σ*_*b*_ and *σ*_*ξ*_ respectively. In addition to background and signal noise, channel current 𝓘 is also treated as normal random variable of mean < 𝓘 > and deviation *σ*_*ξ*_. n_*t*_ is the number of channels open at time t, which is 0 or 1 for current case but can be higher for multi-channel/multi-conductance levels synthetic data.

To test the robustness and accuracy of the software towards noise levels present in the experimental data, we generated single channel synthetic patch-clamp data with SNR ranging from 2.1 to 12.6 as described below.

As mentioned above, there are two noise sources in our model, *ξ*_*t*_ and 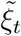:

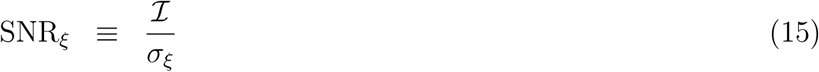

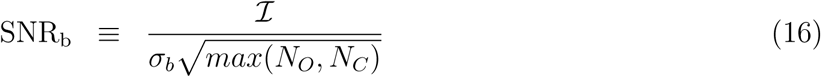

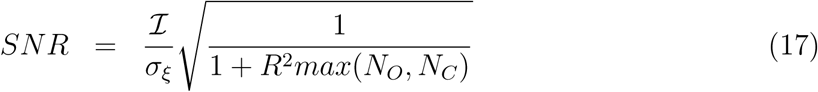

where *N*_*O*_ and *N*_*C*_ are the mean number of sampling intervals for open and closed event respectively. SNR_*ξ*_ is a standard signal to noise ratio. SNR_b_ indicates how much the baseline drifts relative to the current during the average open or closed event. *SNR* is the combined signal to noise ratio: SNR 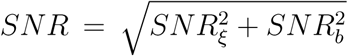 Using 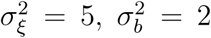, we vary mean current 𝓘 from 20 to 120 (arbitrary current units) to get a SNR from 2.1 to 12.6.

We also process fluorescence data from a simulated cluster composed of 10 IP_3_R Ca^2+^ channels. Details about simulating a single cluster have been published before (26, 29, 30). Briefly, channels were arranged in a 2D array on a patch of endoplasmic reticulum (ER) membrane with an inter-channel spacing of 120nm. The gating of each channel is given by a four state Markov chain model with rest (**R**), active (**A**), open (**O**), and inactivate (**I**) states connected in a loop (Fig. 2 of Ref.(26)). The channel is open when in state **O** and closed otherwise. The state of each channel was determined by using a stochastic scheme outlined in (31).

**Fig. 2:**
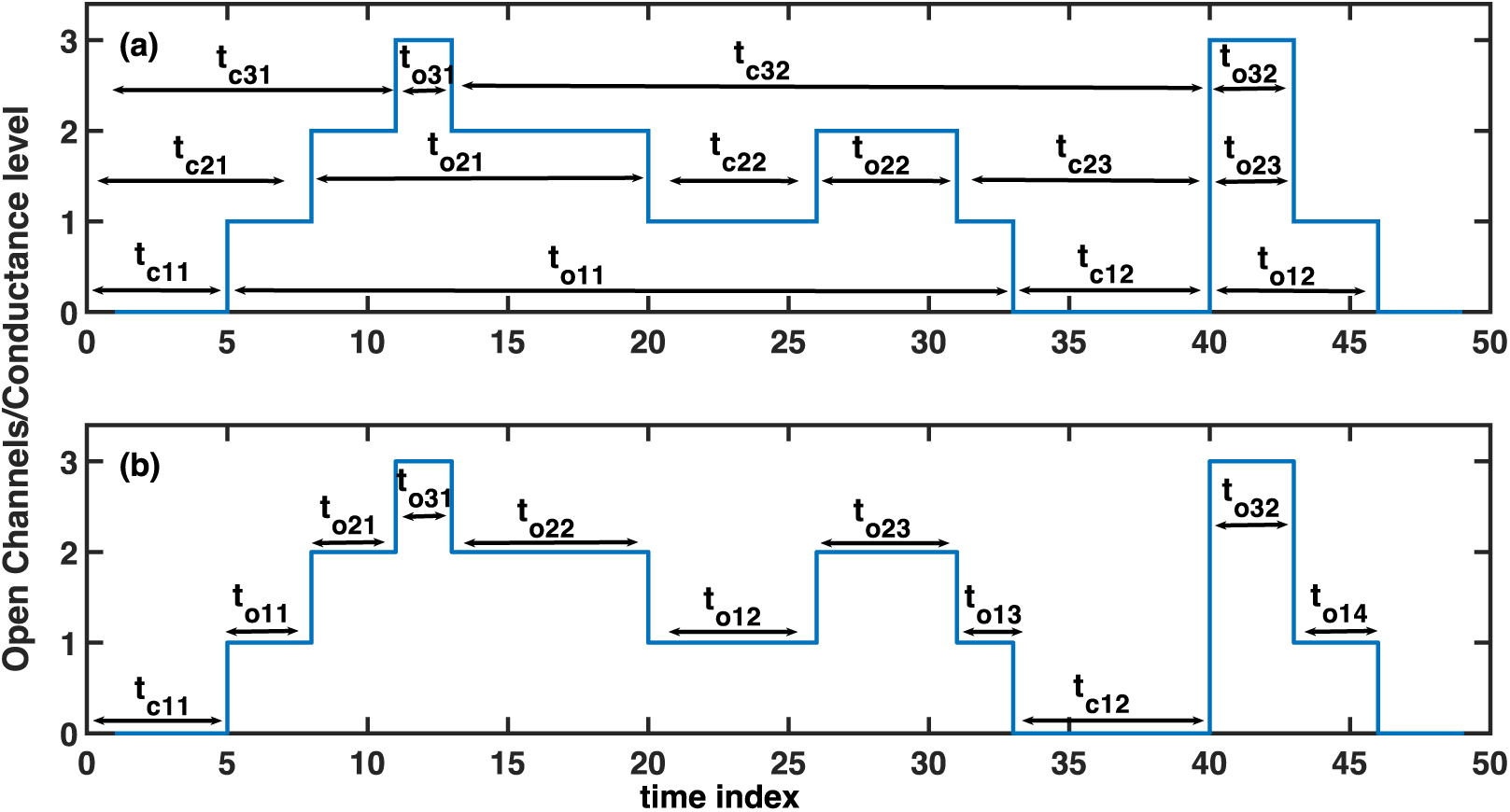
TraceSpecks uses two schemes for calculating open and close dwell-times. As an example, we show an idealized trace with three channels/open conductance levels. (a) In scheme 1, t_oij_ and t_cij_ represent the j^th^ dwell-time when a minimum of i channels are open (or the channel is gating in conductance level i or above) and the j^th^ dwell-time when a minimum of i channels are not open respectively. (b) In scheme 2, t_oij_ represents the j^th^ dwell-time when i channels are open (or the channel is gating in conductance level i) and t_clj_ represents the j^th^ dwell-time when no channel is open (or the channel is in closed level).

Ca^2+^ concentration on the cytosolic side of the cluster was controlled by diffusion, flux through IP_3_Rs, free stationary buffers, mobile buffers, and imaging dye. We considered slow mobile buffer mimicking ethylene glycol tetraacetic acid (EGTA). The propagation of Ca^2+^ and buffers was solved implicitly using the Laplacian of Ca^2+^ and buffers in spherical coordinates on a hemispherical volume of radius 5*μm* and spatial grid size of *5nm*. We estimated the fluorescence changes (Δ*F*/*F*_0_) from TIRF microscopy by following the procedure in (32, 33), i.e.

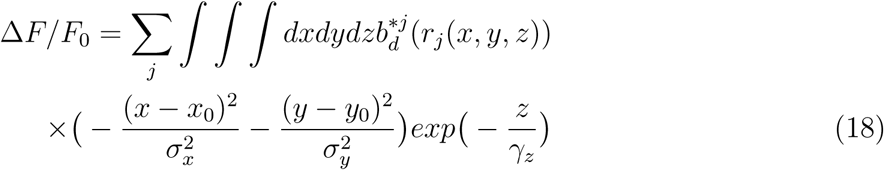

Where 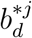 is the Ca^2+^-bound dye at distance *r*_*j*_(*x,y,z*) due to Ca^2+^ released by the *j*^*th*^ channel at *x*_0_ = 0, *y*_0_ = 0, 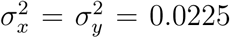 and *γ*_*z*_ = 0.15 *μm.* The integration in Eq. 18 was performed over a cubic volume of 1.5 *μm* × 1.5 *μm* × 0.15 *μm*. To make the simulated TIRF signal as close to the experiment as possible, we added experimental noise and drifting background extracted from TIRF microscopy of puff sites in Xenopus oocytes to the simulated fluorescence data.

#### Experimental data sets

Experimental traces of current passing through single IP_3_R channels, which are ubiquitous intracellular Ca^2+^-release channels localized mainly to the ER and outer nuclear membranes (34), were acquired by nuclear patch-clamp electrophysiology as described in Refs.(27, 35, 36). Currents passing through outer nuclear membrane patches isolated at the tips of micro-pipettes were amplified using an Axopatch 200B patch-clamp amplifier (Molecular Devices, Sunnyvale, CA); filtered either at 1 kHz using a tunable low-pass four-pole Bessel filter (Frequency Devices, Ottawa, IL) or at 5 or 10 KHz using the internal tunable low-pass four-pole Bessel filter of the Axopatch amplifier. The current signals were digitized at 5 kHz using an ITC16 interface (HEKA Instruments, Bellmore, NY) and recorded directly onto a data acquisition computer using the Pulse + PulseFit software (HEKA Instruments, Bellmore, NY).

Full details of TIRFM experiments on IP_3_-induced puffs are given in Ref.(28). Briefly, records of IP_3_-induced puffs were obtained from Ca^2+^ imaging of Xenopus laevis oocytes at room temperature by widefield epifluorescence microscopy using an Olympus inverted microscope (IX 71) equipped with a 60xoil-immersion objective, a 488 nm argon ion laser for fluorescence excitation, and a CCD camera (Cascade 128+; Roper Scientific) for imaging fluorescence emission (510 - 600 nm) at frame rates of 30 to 100 *s*^−1^. Fluorescence was imaged within a 40 × 40 *μ*m region within the animal hemisphere of the oocyte, and measurements are expressed as a ratio (ΔF/F_0_) of the mean change in fluorescence (ΔF) at a given region of interest (1.5 *μm* × 1.5 *μm*) centered on the putative IP_3_R cluster, relative to the resting fluorescence at that region before stimulation (F_0_). Mean values of F_0_ were obtained by averaging over several frames before stimulation. MetaMorph (Molecular Devices) was used for preliminary image processing. Fluorescence traces representing Ca^2+^ release events due to channels’ gating in the cluster were extracted from the image sequences using a previously developed software called CellSpecks that can detect all Ca^2+^ events in a video sequence recorded from cell’s membrane and extracts fluorescence time-traces for individual clusters. Further details about CellSpecks can be found in Ref. (8).

Detailed experimental methods for imaging the activity of PM pores formed by A*β*_1-42_ oligomers are given in Ref.(8). Briefly, solution containing soluble oligomers prepared from human recombinant A*β*1-42 peptide and aliquots were applied using a glass pipette of tip diameter of ≈ 30 *μ*m to voltage-clamped oocytes of defolliculated stage VI Xenopus treated with fluo-4 dextran. For imaging, oocytes were placed animal hemisphere down in a chamber whose bottom is formed by a fresh ethanol washed microscope cover glass (type-545-M; Thermo Fisher Scientific) and were bathed in Ringer solution (110 mM NaCl, 1.8 mM CaCl2, 2 mM KCl, and 5 mM Hepes, pH 7.2) at room temperature (≈23°C) continually exchanged at a rate of ≈ 0.5 ml/min by a gravity-fed superfusion system. The membrane potential was clamped at a holding potential of 0 mV using a two-electrode voltage clamp (Gene Clamp 500; Molecular Devices) and was stepped to more negative potentials −100 mV when imaging Ca^2+^ flux through amyloid pores to increase the driving force for Ca^2+^ entry into the cytosol. The video sequences generated by TIRFM were processed by CellSpecks to extract fluorescence traces for individual A*β*_1-42_ pores.

## Results and Discussion

We use TraceSpecks to idealize experimental time traces from patch-clamp experiments with single and multiple channels in the patch, TIRFM of Ca^2+^ puffs generated by clusters of IP_3_Rs, fluorescence traces representing the gating of Ca^2+^- permeable pores formed by A*β*_1-42_ oligomers in the PM, simulated TIRF signals from a cluster of IP_3_R channels as well as synthetic patch-clamp data with variable SNR. In this section, we first demonstrate the robustness and accuracy of our software using synthetic data with variable SNR and then apply it to other data sets mentioned above.

Before presenting our results, we would like to point out that the software uses two different schemes for reporting the dwell-time distributions, mean P_*O*_, *τ*_*O*_, and *τ*_*C*_ for different conductance levels or number of open channels in the trace as shown in Fig. 2. Although we only present the dwell time distributions and other statistics from the first scheme (Fig. 2a) in the paper, TraceSpecks saves the results from both schemes in ascii format in separate files.

### Synthetic data

As described in the “Methods” section, using 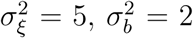, and varying mean current 𝓘 from 20 to 120 (arbitrary units), we have generated synthetic data sets with SNR ranging from 2.1 to 12.6. As shown in Fig. 3, TraceSpecks idealized these noisy synthetic patch-clamp traces with drifting and variable background, and accurately determined various parameters of interest for almost all values of SNR. A sample noisy trace with the background separated by the software and the idealized trace are shown in Fig. 3a, and c respectively. We also show the actual trace without the noise and background for comparison in Fig. 3b. We repeat this experiment for many traces with varying SNR, estimate different features, and compare these estimates with the actual values of these features. In Fig. 3d-f we show the mean squared error for *τ*_*O*_, *τ*_*C*_, and P_*O*_. Even for SNR as small as 2.1, the estimates for these parameters are very good. It is worth mentioning that the SNR in TIRFM experiments is close to 8 (37) while that for the patch-clamp electrophysiology is even higher (38). TraceSpecks extracts the exact values for different parameters of interest for traces with SNR of 5 or above (Fig. 3d-f).

**Fig. 3:**
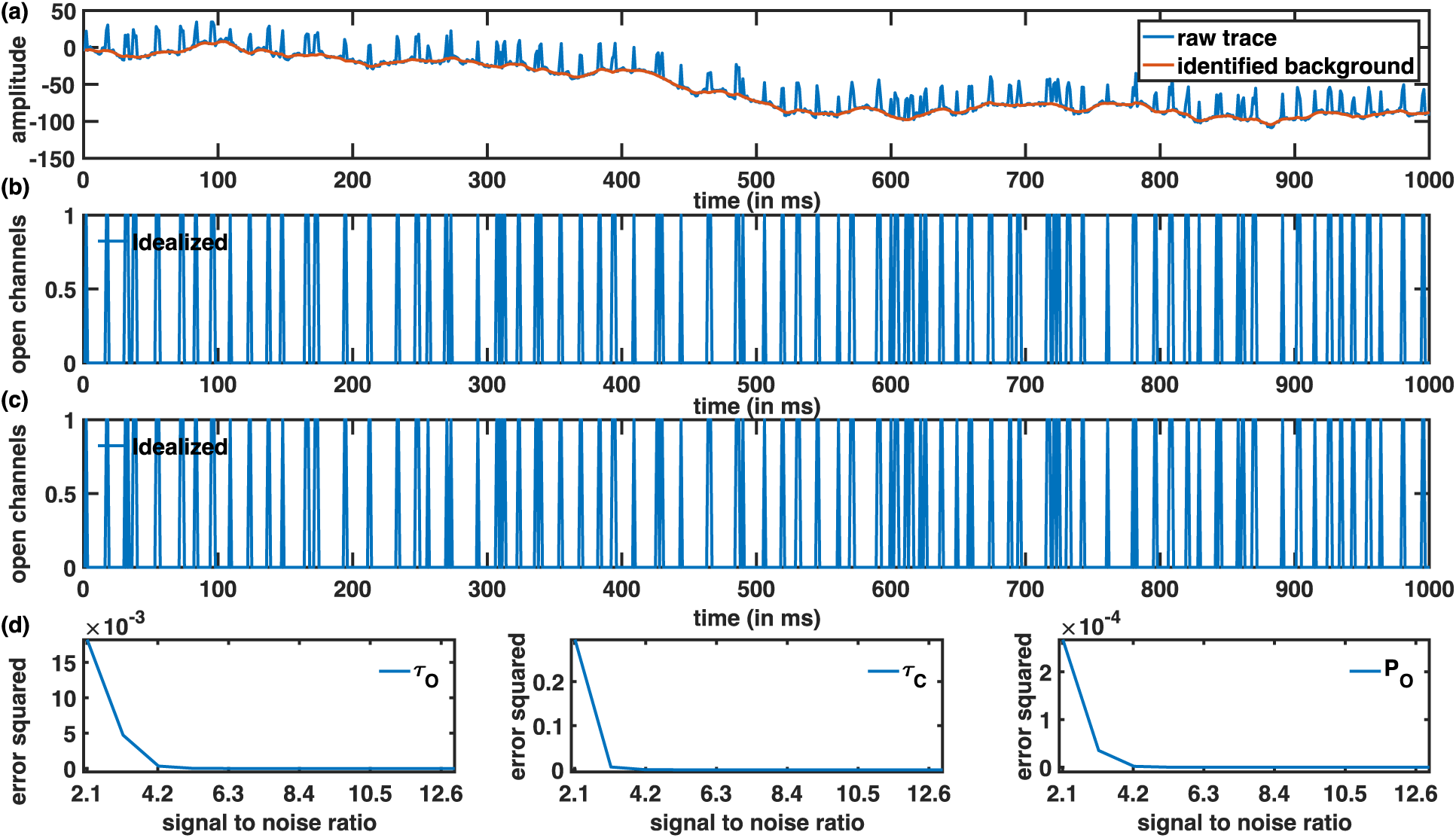
Processing synthetic patch-clamp data with variable SNR of 2.1 – 12.6. (a) Sample raw trace with noisy and drifting background (blue line) and background separated by TraceSpecks (red line), (b) actual trace, and (c) idealized trace. Mean squared error for mean *τ*_*O*_ (d), mean *τ*_*C*_ (e), and mean P_*O*_ (f).

Application to fluorescence data from a simulated cluster, composed of 10 IP_3_R channels demonstrates (black line in Fig. 4a.) that the software works well for multi-channel data too. The drifting background separated from the signal by the software is shown by the red line in Fig. 4a. The idealized signal representing the expected number of open channels in the cluster at time *t* is shown in Fig. 4b. For simulations shown in Fig. 4a, we also recorded the number of open channels in the cluster that is shown in Fig. 4c for comparison. It is clear from Fig. 4b and c that the processed signal closely compares with the actual number of open channels. It should be noted that our method also reports *τ*_*O*_, *τ*_*C*_, P_*O*_ as well as the dwell-time distributions for different number of open channels and inter-channel transitions (transition from X number of channels open to Y number of channels open at a time t where Y can be less than or larger than X) for the given trace (not shown for this example).

**Fig. 4:**
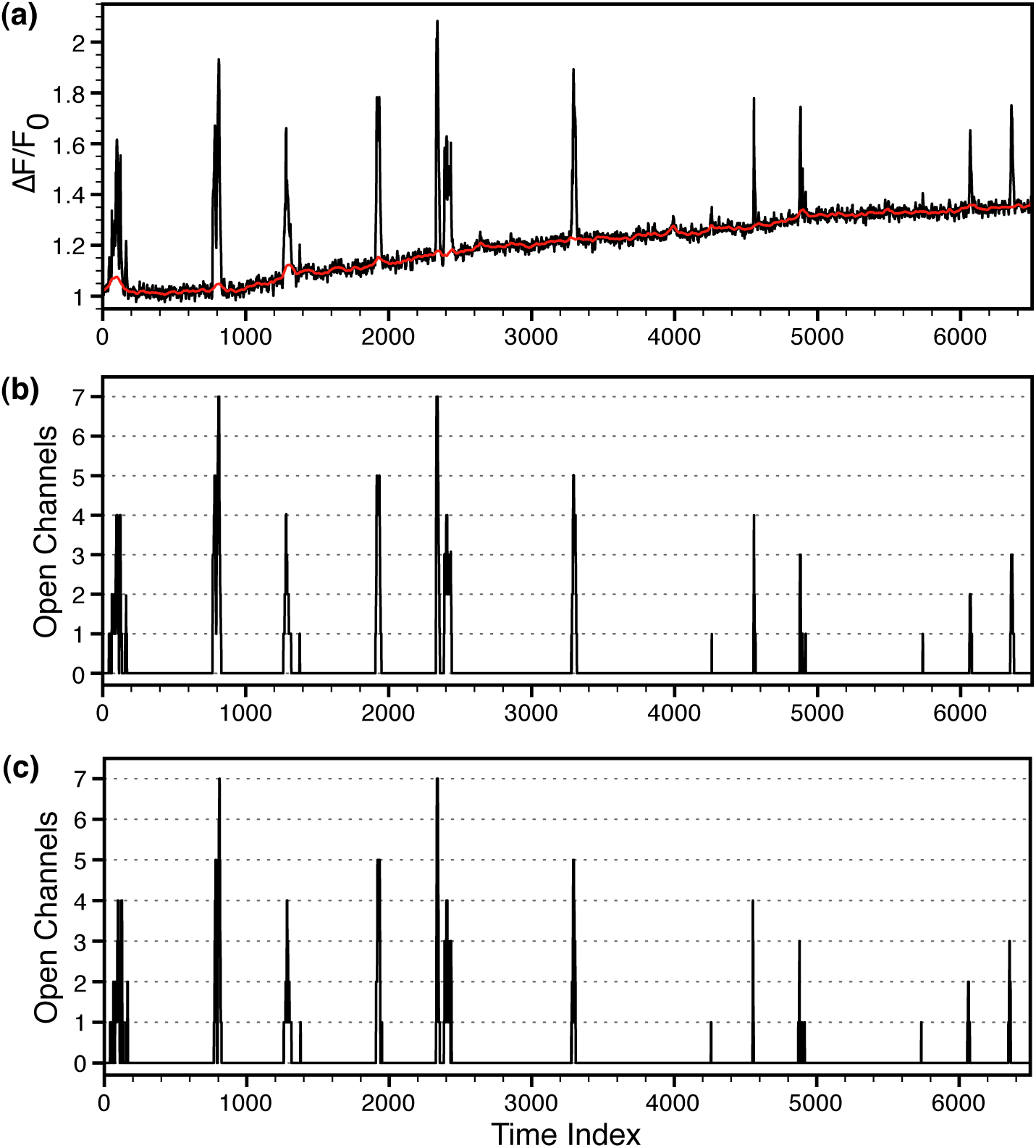
Processing simulated puffs. (a) Noisy and drifting fluorescence time-series generated by a simulated single IP_3_R cluster with added background extracted from the raw experimental data (black) and the drifting background estimated by the software (red). (b) Idealized signal generated by the software. (c) Number of open channels in the cluster as a function of time from simulation shown for comparison.

### Experimental data

In this section, we demonstrate the application of our method to fluorescence time-traces representing IP_3_-induced Ca^2+^ puffs by IP_3_R clusters in the ER membrane and the gating of PM pores formed by Α*β*_1-42_ oligomers, both observed through TIRFM in Xenopus oocytes, as well as traces from nuclear patch-clamp electrophysiology of IP_3_Rs in Sf9 cells with single and multiple channels per patch.

First, we idealize traces from nuclear patch-clamp electrophysiology of IP_3_Rs. While the method works equally well for traces with single channel per patch (see for example Fig. 3), we only show slightly complex examples of multiple channels per patch. A summary of eight traces with two to four channels per patch is shown in Fig. 5 and Table 1. A sample raw trace with two channels along with the separated background and idealized trace identified by the algorithm is shown in Fig. 5a and b respectively. Dwell-time distributions when at least one, two, three, or four channels are open in all eight traces are shown in Fig. 5c-e respectively. Our method also provides mean P_*Oi*_, *τ*_*Oi*_, and *τ*_*Ci*_ for at least 𝓘 channels open for the eight traces (Table 1). For example, P_*O*1_ and *τ*_*O*1_ is the mean probability and mean open time that at least one channel is open. Similarly, *τ_c_*_1_ is the mean time when none of the channels are open (a minimum of one channel is not open) and *τ*_*C*2_ is the mean time when either one or no channels are open (a minimum of two channels are not open).

**Table 1:** Channel statistics for eight traces from nuclear patch-clamp electrophysiology of IP_3_Rs in Sf9 cells with two (trace 1-3), three (trace 4-6), and four (trace 7 and 8) active channels per patch. Columns 2-4 and 5-8 are the mean probabilities and mean open times respectively when at least one, two, three, and four channels are open. Columns 9-12 are the mean close times when a minimum of one, two, three, and four channels are not open.

| Trace | $P_{O1}$ | $P_{O2}$ | $P_{O3}$ | $P_{O4}$ | $\tau_{O1}$<br>*ms* | $\tau_{O2}$<br>*ms* | $\tau_{O3}$<br>*ms* | $\tau_{O4}$<br>*ms* | $\tau_{C1}$<br>*ms* | $\tau_{C2}$<br>*ms* | $\tau_{C3}$<br>*ms* | $\tau_{C4}$<br>*ms* |
| --- | --- | --- | --- | --- | --- | --- | --- | --- | --- | --- | --- | --- |
| 1 | 0.91 | 0.67 | - | - | 166 | 271 | - | - | 17 | 128 | - | - |
| 2 | 0.41 | 0.16 | - | - | 642 | 163 | - | - | 927 | 8017 | - | - |
| 3 | 0.82 | 0.52 | - | - | 505 | 790 | - | - | 111 | 722 | - | - |
| 4 | 0.93 | 0.86 | 0.38 | - | 113 | 138 | 171 | - | 9 | 23 | 278 | - |
| 5 | 0.41 | 0.14 | 0.003 | - | 187 | 275 | 160 | - | 265 | 1683 | 36525 | - |
| 6 | 0.36 | 0.14 | 0.02 | - | 29 | 53 | 75 | - | 51 | 321 | 3558 | - |
| 7 | 0.69 | 0.41 | 0.11 | 0.002 | 49 | 72 | 91 | 77 | 22 | 105 | 714 | 21639 |
| 8 | 0.81 | 0.60 | 0.30 | 0.002 | 911 | 1566 | 2166 | 1100 | 219 | 1048 | 4922 | 253520 |

**Fig. 5:**
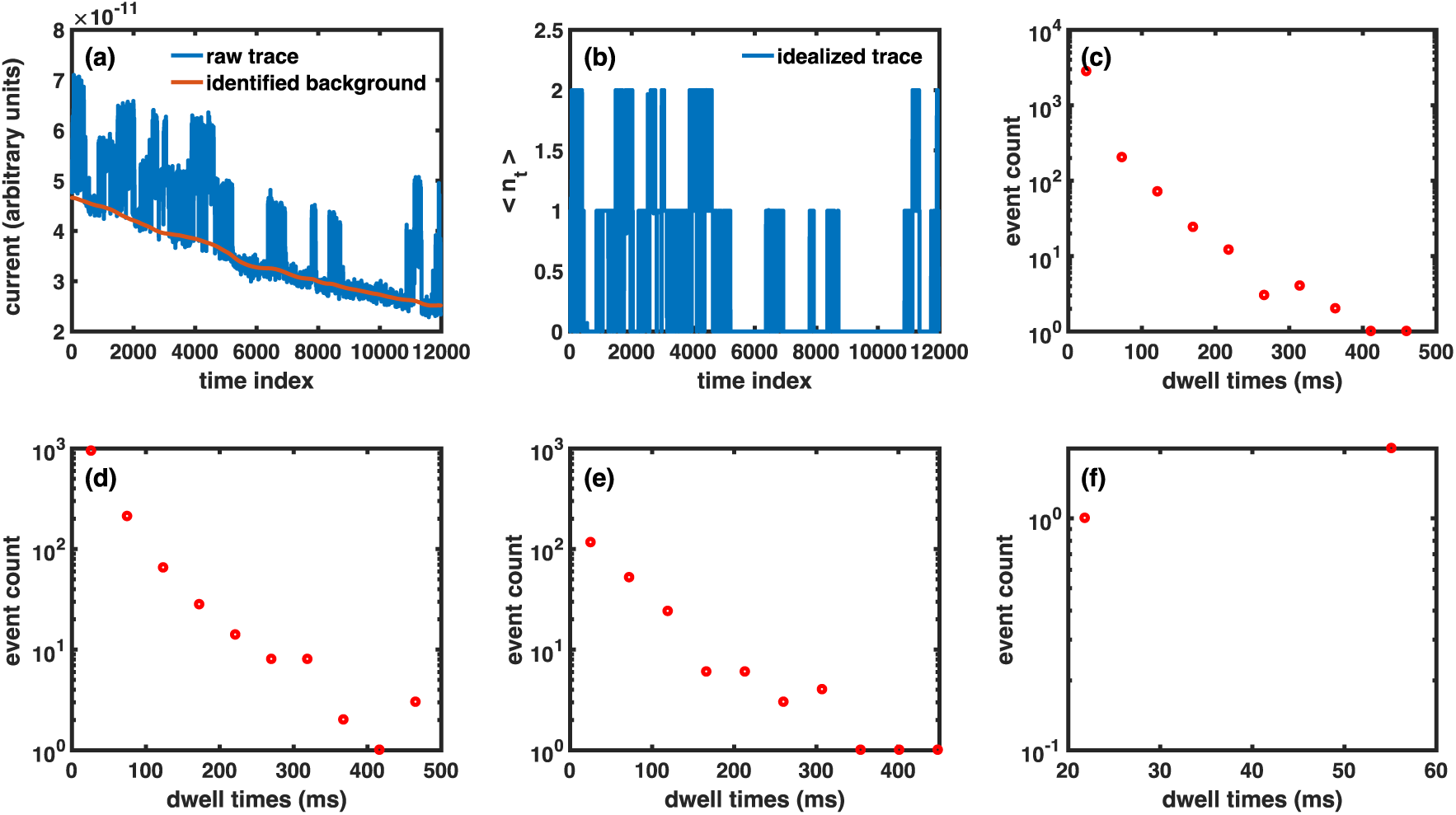
Processing current time-traces that record the gating of IP_3_R channels during single-channel patch-clamp electrophysiological experiments using nuclei isolated from Sf9 cells. (a) A typical raw current trace (blue line) in which two channels were active, and the background leak current level (red) identified by our software. (b) The number of open channels at any time during the experiment in (a). Dwell-time distributions when at least one (c), two (d), three (e), or four channels are open in all eight traces processed.

We also processed many fluorescence records from TIRFM of IP_3_-induced Ca^2+^ puffs in Xenopus oocytes with varying amplitudes (in terms of peak fluorescence or number of open channels during the puff) and duration. We identified up to ten channels in the traces processed and show a range of parameters and statistics extracted from the fluorescence traces by our software in Fig. 6. Panels (a1) and (a2) show a raw trace, representative of IP_3_-induced puffs (blue) with background noise (red) and the number of channels identified by TraceSpecks respectively. The raw TIRFM signal is the relative fluorescence change due to the cluster activity, averaged over a 1.5 *μm* × 1.5 *μm* area surrounding the cluster. As pointed out above, the software allows us to store the mean P_*O*_, *τ*_*O*_, *τ*_*C*_, and dwell-time distributions for different number of channels open in the cluster, and amplitudes of all puffs in a trace in ascii format for plotting and post-processing. As an example, we show the distribution of puff amplitudes represented as the maximum number of channels opened simultaneously in a puff from all traces processed in panel (a3) of Fig. 6.

**Fig. 6:**
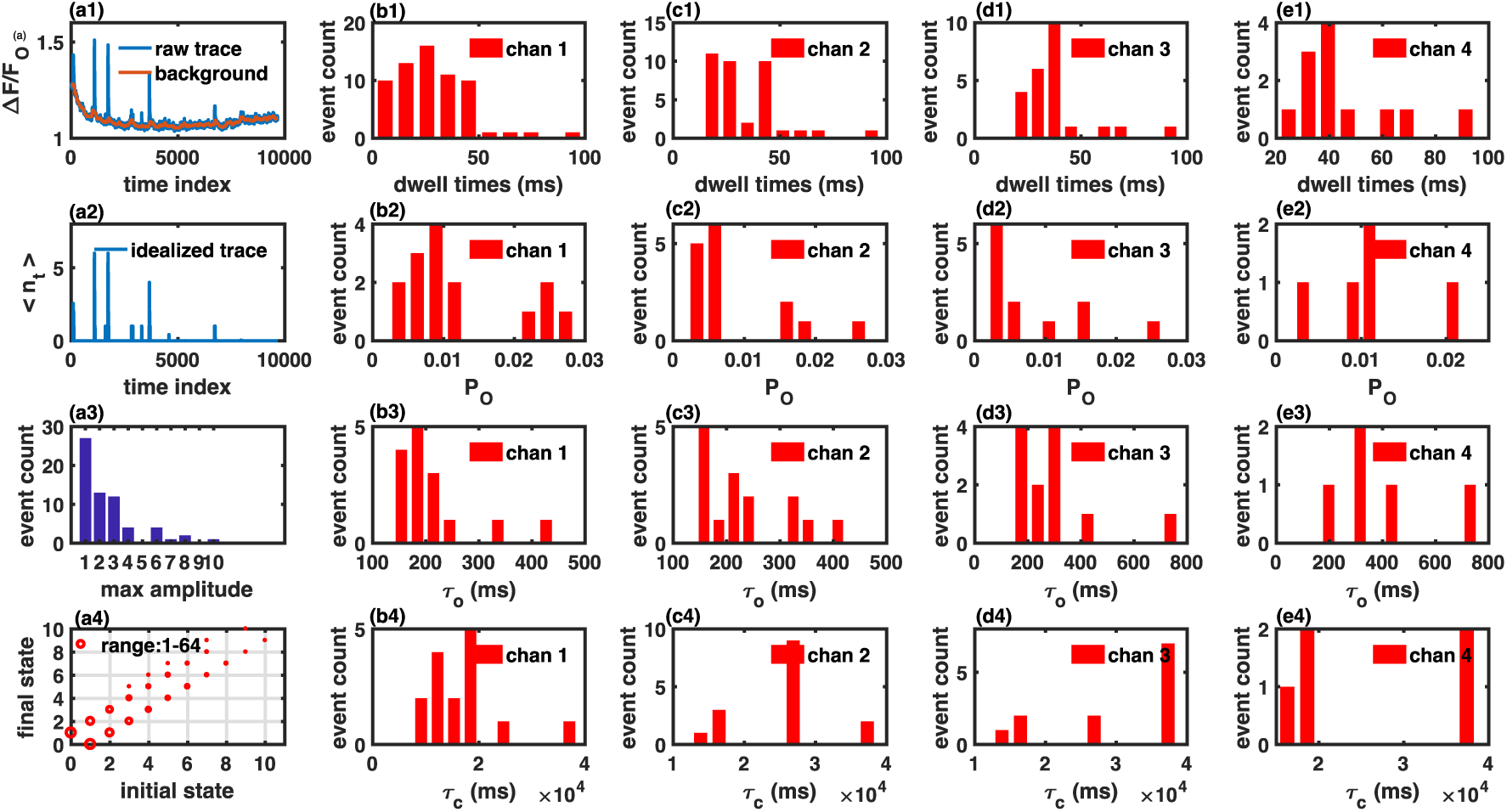
Processing fluorescence records from TIRFM of IP3-induced Ca^2+^ puffs in Xenopus oocytes. The first column on the left shows a sample raw trace (blue) and the background separated by the software (red) (a1), the corresponding idealized trace (a2), amplitude (maximum number of channels open simultaneously) distribution of all puffs in all traces processed, and the transition between different number of channels open/close (a4). The remaining four columns show distributions of dwell-times (first row), mean P_*O*_ (second row), mean *τ*_*O*_ (third row), and mean *τ*_*C*_ (fourth row) when at least one (panels b1-b4), two (panels c1 - c4), three (panels d1 - d4), and four channels (panels e1 - e4) are open in all traces processed. Mean *δ*_*C*_ for channel *i* represents the mean time when the number of open channels is less than *i*.

Another key information about the traces representing the activity of multiple channels or single channel with multiple conductance levels is determining the probability of transitions between different number of channels opened or different conducting states. For example, what is the likelihood that Y channels will be open at time t+*δ*t given that there are X channels open at time t, where X and Y don’t have to be consecutive digits? Or what is the probability that a channel in conductance level X at time t will be in conductance level Y at time t+*δ*t? This information can be used for modeling the kinetics of a single channel with multiple conductance levels or a cluster with multiple channels. We report this information as an inter-state or inter-channel transition graph as shown in panel (a4). In this graph, the point at location (X,Y) = (2,1) indicates that there is a transition from 2 simultaneous open channels to 1 open channel (one of the open channels closed). The size of the circle is proportional to the number of such transitions found in all traces we processed. Similarly, the point at location (X,Y) = (1,2) indicates that another channel opened up between time t and t+*δ*t and the total number of open channels is now 2. We also observed transitions where the total number of channels at time t and t+*δ*t are not consecutive digits. The dwell-time distributions (first row), mean P_*O*_ (second row), mean *τ*_*O*_ (third row), and mean *τ*_*C*_ (fourth row) when at least one (panels b1 - b4), two (panels c1 - c4), three (panels d1 - d4), and four (panels e1 - e4) channels are open in all traces are also shown in Fig. 6. We remark that the software saves this information for all channels identified (ten in this case) but we skip the statistics for the remaining six channels for clarity.

The method also works for fluorescence traces representing the function of ion channels having multiple levels with quantal conductances and traces with very low SNR. As an example, we process fluorescence traces from TIRFM of many A*β*_1-42_-induced Ca^2+^-permeable pores in the PM of Xenopus oocyte and summarize our results in Fig. 7. We observe a closed level where no Ca^2+^ flux is passing through the pore (represented by level 0) and up to five open conductance levels (represented by level 1, 2, 3, 4, and 5) (9). In Fig. 7, we show a sample raw trace, the separated background, and idealized trace, the distribution of dwell times, mean P_*O*_, mean *τ*_*O*_, mean *τ*_*C*_ when the channel is gating in a conductance level equal to or larger than 1, 2, 3, and 4, and the number of transitions between different conductance levels. We skip the distributions for conductance level 5 as there were only a few instances where the channel opened in that level. The details about the data for A*β*_1-42_ pores is similar to that of IP_3_-induced puffs except that the number of open channels from puffs discussion is replaced by conducting levels in case of A*㭂*_1-42_ pores. We would like to point out that there are significantly larger number of transitions between levels 0 and 1 in comparison to the puffs data, indicating that A*β*_1-42_ pores prefer to gate in low conductance levels. Also, in comparison to IP_3_-induced puffs, we observed a larger number of transitions where the levels involved are not immediately next to each other (for example transitions from level 0 to level 2 and vice versa) (Fig. 7a4).

**Fig. 7:**
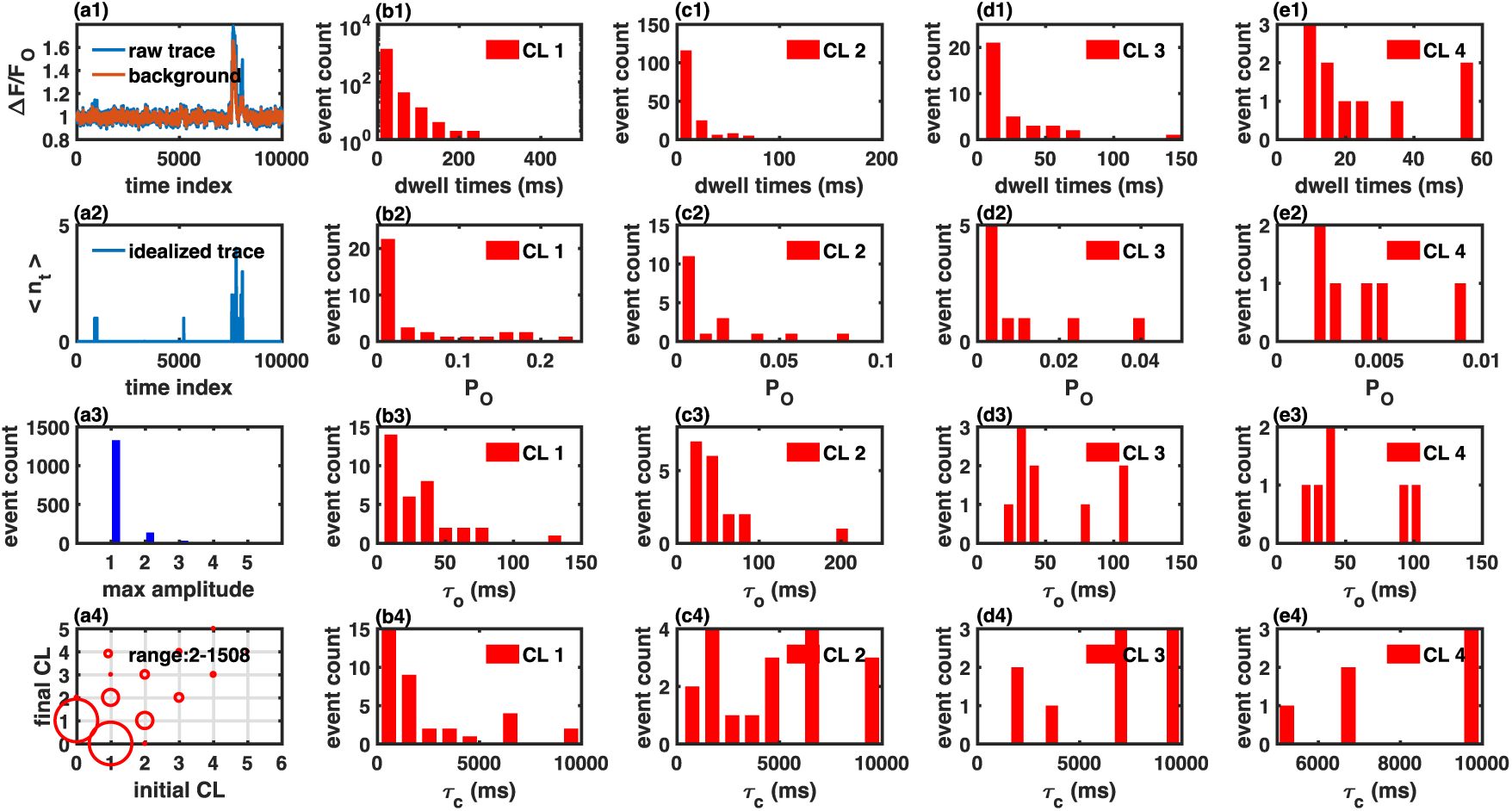
Processing fluorescence records from TIRFM of A*β*_1-42_-induced pores in the PM of Xenopus oocytes. The first column on the left shows a sample raw trace (blue) and the background separated by TraceSpecks (red) (a1), the corresponding idealized trace representing the conductance level in which the pore is gating (a2), amplitude (the highest conductance level in which the channel is gating during a single opening event) distribution, and the transition between different conductance levels (a4) for all opening and closing events in all traces processed. The remaining four columns show distributions of dwell-times (first row), mean P_*O*_ (second row), mean *τ*_*O*_ (third row), and mean *τ*_*C*_ (fourth row) when the pore is gating in level 1 or above (panels b1-b4), level 2 or above (panels c1 - c4), level 3 or above (panels d1 - d4), and level 4 or above (panels e1 - e4) in all traces processed. Mean *τ*_*C*_ for conductance level i represents the mean time when the conductance level is less than *i*.

## Conclusions

Data from fluorescence imaging as well as the routinely employed electrophysiological patch-clamp techniques are contaminated with noise and drifting background that does not arise from the system under study. Removing this noise and fluctuating background is usually the first step in the analysis of the data. In the absence of automated method, experimentalists adhere to the eye and mouse-based approach for processing noisy imaging and patch-clamp data. This laborious manual procedure could cost experimentalists more time for processing the data than the actual experiment.

One example of where the lack of efficient algorithm for separating the signal from background noise could limit the full utilization of very powerful experimental techniques is the high resolution imaging of Ca^2+^ signals. Recent advances in imaging techniques such as TIRFM offer a paradigm shifting improvement in our ability to study thousands of ion channels in parallel in their native environment (1, 2). This technique called “optical patch-clamp” generates fluorescence time traces from thousands of ion channels in a single experiment. While an algorithm to automatically detect, localize, and measure fluorescence changes during localized Ca^2+^ signals, imaged through optical patch-clamp has been recently developed (39), the lack of efficient algorithms to extract signal out of the noise and idealize the fluorescence traces remains the main hurdle in accessing a wide range of information about the system under investigation and developing single channel/molecule models based on huge data-sets collected from thousands of channels/molecules in their native environment in these experiments.

In this work, we developed a minimally-parameterized likelihood method that can process both noisy patch-clamp and imaging data with drifting background. The robustness and accuracy of our algorithm is demonstrated by closely recovering the signal from noisy synthetic patch-clamp data with variable SNR and fluorescence traces representing Ca^2+^ puffs generated by a cluster of IP_3_R channels. This was followed by the processing of significant data from both TIRFM experiments as well as patch-clamp recordings and extracting many important features from the data.

We implemented our algorithm in the very portable java language with options to setup initial parameters, display optimized parameters, raw trace, separated background, and idealize trace, mean P_*O*_, *τ*_*O*_, and *τ*_*C*_ for all channels/conductance levels in the trace, and overall mean P_*O*_, *τ*_*O*_, and *τ*_*C*_ of the trace (Fig 1). Option to store all these traces and statistics in addition to dwell-times for the channel gating in different conducting states or different number of open channels in a cluster or a patch, amplitude distribution of all events in terms of the highest number of channels open or conductance level in which the channel is gating during the event as well as the number of transitions between different states in ascii format for plotting and later use is also provided. A compiled java program with user-friendly GUI that performs all these tasks, portable to Mac, PC, and Unix platforms with user manual, will be provided in the online supplement upon acceptance of this paper for publication.

## Acknowledgements

This work was supported by NIH through grants R01 AG053988 (to AD and GU), R01 GM065830 (to DODM, IP, and JEP), and R37 GM048071 (to IP).

## References

1. Demuro, A., and I. Parker, 2004. Imaging the activity and localization of single voltage-gated Ca^2+^ channels by total internal reflection fluorescence microscopy. Biophys. J. 86:3250–3259.

2. Demuro, A., and I. Parker, 2005. Optical patch-clamping: single-channel recording by imaging Ca^2+^ flux through individual muscle acetylcholine receptor channels. J. Gen. Physiol. 126:179–192.

3. Demuro, A., and I. Parker, 2004. Imaging single-channel calcium microdomains by total internal reflection microscopy. Biological Research 37:675–679.

4. Demuro, A., and I. Parker, 2005. Optical single-channel recording: imaging Ca^2+^ flux through individual ion channels with high temporal and spatial resolution. Journal of Biomedical Optics 10:011002–0110028.

5. Demuro, A., and I. Parker, 2006. Imaging single-channel calcium microdomains. Cell Calcium 40:413–422.

6. Smith, I., and I. Parker, 2009. Imaging the quantal substructure of single IP_3_R channel activity during Ca^2+^ puffs in intact mammalian cells. Proc. Natl. Acad. Sci. 106:6404–6409.

7. Navedo, M. F., and L. F. Santana, 2013. CaV1. 2 sparklets in heart and vascular smooth muscle. Journal of molecular and cellular cardiology 58:67–76.

8. Demuro, A., M. Smith, and I. Parker, 2011. Single-channel Ca^2+^ imaging implicates A*β*1-42 amyloid pores in Alzheimers disease pathology. J. Cell Biol. 195:515–524.

9. Ullah, G., A. Demuro, I. Parker, and J. E. Pearson, 2015. Analyzing and modeling the kinetics of amyloid beta pores associated with Alzheimers disease pathology. PloS one 10:e0137357.

10. Colquhoun, D., 1987. Practical analysis of single channel records. *In* Microelectrode techniques: the plymouth workshop handbook, Company of Biologists, 101–139.

11. Witkoskie, J. B., and J. Cao, 2008. Analysis of the Entire Sequence of a Single Photon Experiment on a Flavin Protein. The Journal of Physical Chemistry B 112:5988–5996.

12. Sachs, F., 2009. Automated Analysis of Single-Channel Records. In Single-channel recording, Springer.

13. Colquhoun, D., and F. Sigworth, 2009. Fitting and Statistical Analysis of Single-Channel Data. In Single-channel recording, Springer.

14. Milescu, L., A. Yildiz, P. Selvin, and F. Sachs, 2006. Extracting dwell time sequences from processive molecular motor data. Biophysical Journal 91:3135–3150.

15. Milescu, L., 2003. Applications of hidden Markov models to single molecule and ensemble data analysis. Ph.D. thesis, SUNY, Buffalo.

16. Hamill, O., A. Marty, E. Neher, B. Sakmann, and F. Sigworth, 1981. Improved patch-clamp techniques for high-resolution current recording from cells and cell-free membrane patches. Pflügers Archiv. Eur. J. Physiol. 391:85–100.

17. Qin, F., A. Milescu, F. Qiong, C. Nicolai, and J. Bannen, 2008. QUB. http://www.qub.buffalo.edu-->.

18. Qin, F., A. Auerbach, and F. Sachs, 1996. Estimating single-channel kinetic parameters from idealized patch-clamp data containing missed events. Biophysical Journal 70:264–280.

19. Colquhoun, D., 2007. HJCFit. http://www.ucl.ac.uk/pharmacology/dcpr95.html#intro-->.

20. Fisher, R., 1925. Theory of statistical estimation. Math. Proc. Cambridge Phil. Soc. 22:700–725.

21. Bruno, W., G. Ullah, D. D. Mak, and J. Pearson, 2013. Automated Maximum Likelihood Separation of Signal from Baseline in Noisy Quantal Data. Biophysical Journal. 105:68–79.

22. Baum, L., and T. Petrie, 1966. Statistical inference for probabilistic functions of finite state Markov chains. The Annals of Mathematical Statistics 37:1554–1563.

23. Baum, L., T. Petrie, G. Soules, and N. Weiss, 1970. A maximization technique occurring in the statistical analysis of probabilistic functions of Markov chains. Annals. Math. Stat. 41:164–171.

24. Welch, L. R., 2003. Hidden Markov models and the Baum-Welch algorithm. IEEE Information Theory Society Newsletter 53:10–13.

25. Dempster, A., N. Laird, and D. Rubin, 1977. Maximum likelihood from incomplete data via the EM algorithm. J. Roy. Stat. Soc. Series. (Methodological) 1–38.

26. Ullah, G., I. Parker, D. Mak, and J. Pearson, 2012. Multi-scale data-driven modeling and observation of calcium puffs. Cell Calcium.

27. Mak, D.-O. D., J. E. Pearson, K. P. C. Loong, S. Datta, M. Fernández-Mongil, and J. K. Foskett, 2007. Rapid ligand-regulated gating kinetics of single inositol 1, 4, 5-trisphosphate receptor Ca^2+^ release channels. EMBO Reports 8:1044–1051.

28. Demuro, A., and I. Parker, 2013. Cytotoxicity of intracellular A*β*42 amyloid oligomers involves Ca^2+^ release from the endoplasmic reticulum by stimulated production of inositol trisphosphate. Journal of Neuroscience 33:3824–3833.

29. Mak, D.-O. D., K.-H. Cheung, P. Toglia, J. K. Foskett, and G. Ullah, 2015. Analyzing and quantifying the gain-of-function enhancement of IP3 receptor gating by familial Alzheimers disease-causing mutants in Presenilins. PLoS Computational Biology 11:e1004529.

30. Ullah, G., and A. Ullah, 2016. Mode switching of Inositol 1, 4, 5-trisphosphate receptor channel shapes the Spatiotemporal scales of Ca^2+^ signals. Journal of Biological Physics 42:507–524.

31. Ullah, G., and P. Jung, 2006. Modeling the statistics of elementary calcium release events. Biophysical Journal 90:3485–3495.

32. Shuai, J., and I. Parker, 2005. Optical single-channel recording by imaging Ca^2+^ flux through individual ion channels: theoretical considerations and limits to resolution. Cell Calcium 37:283–299.

33. Toglia, P. T., G. Ullah, and J. E. Pearson, 2017. Analyzing Optical Imaging of Ca^2+^ Signals via TIRF Microscopy: The Limits on Resolution Due to Chemical Rates and Depth of the Channels. Cell Calcium 67:6573.

34. Foskett, J. K., C. White, K.-H. Cheung, and D.-O. D. Mak, 2007. Inositol trisphosphate receptor Ca^2+^ release channels. Physiological Reviews 87:593–658.

35. Mak, D., S. McBride, and J. Foskett, 1998. Inositol 1, 4, 5-trisphosphate activation of inositol trisphosphate receptor Ca^2+^ channel by ligand tuning of Ca^2+^ inhibition. Proceedings of the National Academy of Sciences of the United States of America 95:15821.

36. Mak, D., C. White, L. Ionescu, and J. Foskett, 2005. Nuclear patch clamp electrophysiology of inositol trisphosphate receptor Ca^2+^ release channels. Methods in Calcium Signaling Research, CRC Press, Boca Raton, FL 203–229.

37. Lock, J. T., I. Parker, and I. F. Smith, 2015. A comparison of fluorescent Ca^2+^ indicators for imaging local Ca^2+^ signals in cultured cells. Cell Calcium 58:638–648.

38. Mak, D.-O. D., H. Vais, K.-H. Cheung, and J. K. Foskett, 2013. Patch-clamp electrophysiology of intracellular Ca^2+^ channels. Cold Spring Harbor Protocols 2013:pdb–top066217.

39. Ellefsen, K. L., B. Settle, I. Parker, and I. F. Smith, 2014. An algorithm for automated detection, localization and measurement of local calcium signals from camera-based imaging. Cell calcium 56:147–156.

